# Sputtered porous Pt for wafer-scale manufacture of low-impedance flexible microelectrodes

**DOI:** 10.1101/2020.02.04.934737

**Authors:** Bo Fan, Alexander V. Rodriguez, Daniel G. Vercosa, Caleb Kemere, Jacob T. Robinson

## Abstract

Recording electrical activity from individual cells in vivo is a key technology for basic neuroscience and has growing clinical applications. To maximize the number of independent recording channels as well as the longevity, and quality of these recordings, researchers often turn to small and flexible electrodes that minimize tissue damage and can isolate signals from individual neurons. One challenge when creating these small electrodes, however, is to maintain a low interfacial impedance by applying a surface coating that is stable in tissue and does not significantly complicate the fabrication process. Here we use a high-pressure Pt sputtering process to create low-impedance electrodes at the wafer scale using standard microfabrication equipment. Direct-sputtered Pt provides a reliable and well-controlled porous coating that reduces the electrode impedance by 5-9 fold compared to flat Pt and is compatible with the microfabrication technologies used to create flexible electrodes. These porous Pt electrodes show reduced thermal noise that matches theoretical predictions. In addition, we show that these electrodes can be implanted into rat cortex, record single unit activity, and be removed all without disrupting the integrity of the coating. We also demonstrate that the shape of the electrode (in addition to the surface area) has a significant effect on the electrode impedance when the feature sizes are on the order of tens of microns. Overall, porous Pt represents a promising method for manufacturing low-impedance electrodes that can be seamlessly integrated into existing processes for producing flexible neural probes.

## 1. Introduction

Microelectrodes for recording neural activity have been used for decades, and are now commonplace for applications including basic neuroscience research^1^ and clinical diagnosis^2^. Traditionally, neural electrodes consist of a rigid backbone (often metal or silicon) that insulated except where exposed conductive sites at as electrodes for sensing electrical activity^3^. Because large and rigid implants cause damage to the soft tissue of the brain and reduce the longevity of recordings^4^, modern electrode designs have prioritized small implant profiles and/or flexible materials that better match the Young’s modulus of brain tissue^5^. Smaller electrodes not only have the advantage of reducing damage to brain tissue, but they also allow for tighter packing of recording sites for a larger coverage of brain tissue and greater discrimination between signals from sources close to each other^6^. Channel counts in modern electrode arrays have increased to the hundreds and thousands as microfabrication technologies have improved^7–8^.

However, in theory, there is a tradeoff between electrode size and signal-to-noise ratio of recording. As the size of electrode sites decrease, total impedance of the electrode will naturally increase: at the interface between the electrode and the brain (also known as the electrode-electrolyte interface), a double layer of polarized ions separates the electrode from the brain, resulting in a capacitive interface^9^. This interface is often modeled as a “Randles cell”, with a capacitor and resistor in parallel representing the double layer capacitance and faradaic resistance of the interface, respectively^9–10^. Others have argued that electrode behavior is better explained by substituting a constant phase element (CPE) in place of a capacitor^11^. In either model, as the surface area of an electrode decreases, so does the capacitance, leading to an increase in impedance. Increased impedance is undesirable in an electrode as it causes an increase in Johnson-Nyquist noise (thermal noise)^10,12^ and greater voltage division of the recorded signal^10^. In developing high-channel count small electrodes, the geometric area of the electrode sites is constrained by the implant size and the desired distance between electrodes, the most common solution is to increase the effective surface area of the electrode sites by adding volumetric conductive polymers^13–14^, rough or porous materials^6,7,15–17^, or materials with 3D topography^18–21^.

There are a number of ways to increase the effective surface area of an electrode, and many are still being discovered and refined. Popular current methods include applying or growing surface coatings such as poly(3,4-ethylene dioxythiophene) (PEDOT)^13–14^, carbon nanorods^16,20^, Pt nanorods^21^, platinum gray^22–24^ or platinum black^6,17^. Alternatively, patterns can be etched via co-sputtering two metals and chemically etching one of them^18,21^, or hydrothermal synthesis of porous material onto a pre-shaped surface^7,25^.

All of these methods are able to reduce the impedance of an electrode, though each requires use of wet chemical processes that are not part of most microfabrication lines and may not be compatible with many types of polymer substrates. Coatings such as PEDOT and platinum black require an electrical connection for electroplating, meaning that electrical hardware and chemical solutions are required to treat the electrodes. This electrochemical deposition step is not a standard process in most commercial microfabrication lines. Co-sputtering metals and dealloying exposes electrodes to harsh, acidic etchants which can damage some types of polymer substrates used for flexible electronics. Another issue with many electrode coatings is that they typically do not covalently bond with the electrode surface making delamination a pervasive issue which requires additional strategies, materials, or processing to mitigate the risk of coating detachment^26, 27^. Flexible substrates further increase the risk of electrode damage or delamination because they undergo greater deformation (i.e. shear or stretch forces) in handling than rigid electrodes.

An ideal method of creating a ‘rough’ electrode would not require additional steps, materials, or equipment and would be compatible with large, wafer-scale production. Here, we introduce a method of direct sputtering a porous platinum coating by altering the parameters of platinum deposition (figure 1(d)). We demonstrate that this method is capable of reducing impedance of electrodes by 5-9 fold, depending on the size of the electrode. In addition, we show that these electrodes are durable enough when applied to a flexible substrate to be electrically tested, implanted, removed, and retested without significantly impacting their impedance.

**Figure 1:**
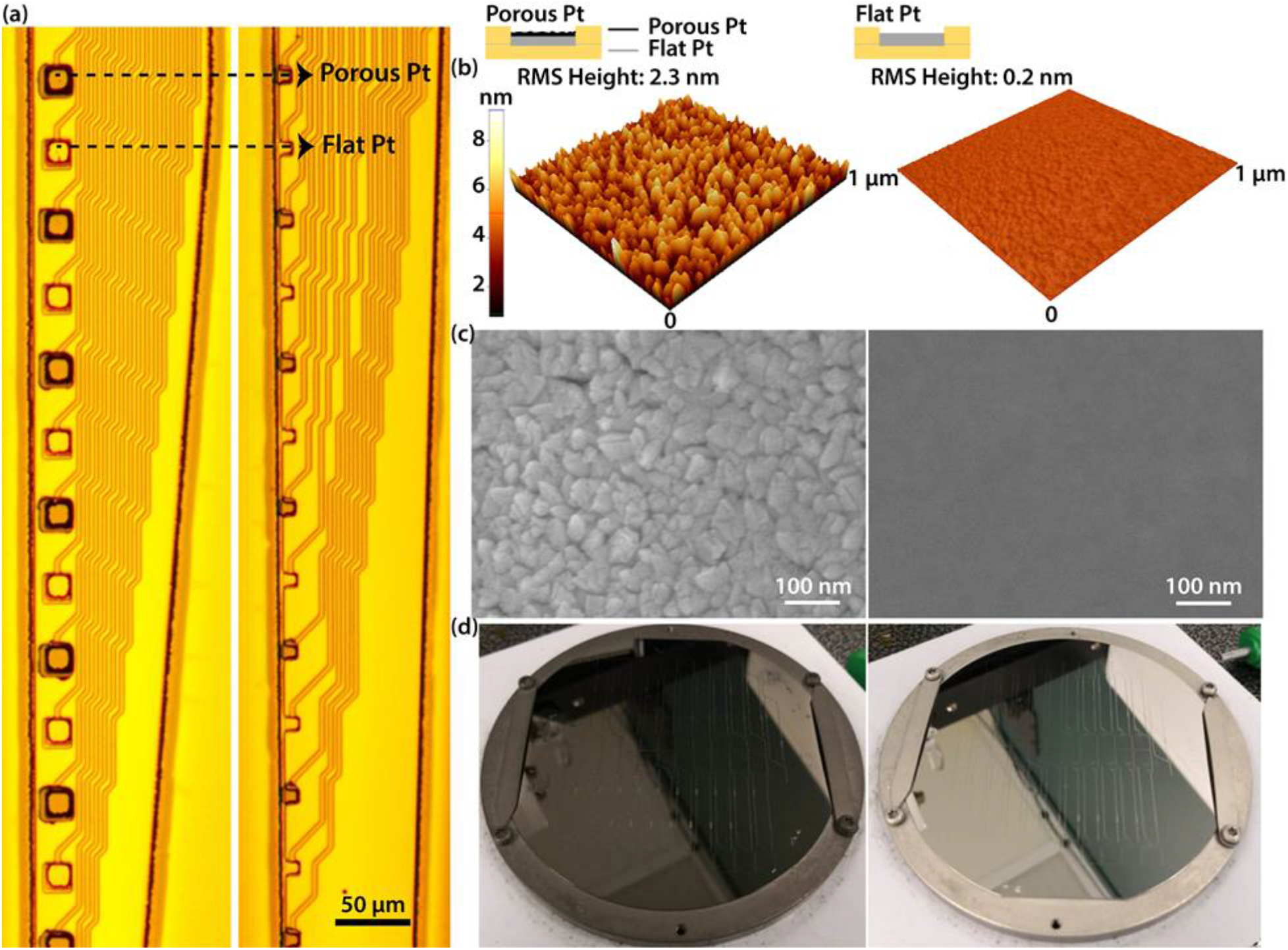
Flexible microelectrode arrays with alternating porous and flat platinum electrodes. **(a)** Two flexible microelectrodes arrays with different sizes of electrodes. 20X microscope images of a probe with 400 μm^2^ electrodes (left) and 100 μm^2^ electrodes (right). Each probe has 16 channels with alternative porous (Dark gray) and flat(Light gray) electrodes. The porous electrodes have a porous platinum layer on top of flat platinum (See inset). Porosity of the Pt layer was determined by changing the sputtering parameters. Scale bar 50 μm. **(b)-(c)** Surface characterization of porous and flat electrodes. Porous platinum layers have larger grain sizes and a rougher surface. **(b)** Atomic force microscope images of a porous (left) and a flat (right) electrode. The root-mean-square (RMS) height of porous Pt and flat Pt are 2.3 nm and 0.2 nm, respectively. The RMS height was calculated using XEI Analysis software 4.3.4, Park Systems Corp. **(c)** Scanning electron microscope images of a porous (left) and a flat (right) electrode. Scale bar 100 nm. **(d)** Directly sputted porous (left) and flat (right) Pt on a 4 inch Si wafer.

## 2. Methods

### 2.1 Fabrication of porous platinum electrodes

We fabricated 16-channel flexible electrodes on an SU-8 substrate using a method similar to that reported previously^28^. Fabrication was performed on a 4-inch silicon wafer (University Wafer). First, a release layer of aluminum roughly 70 nm thick was applied to the entire wafer. Then, a 4-μm-thick layer of SU-8 2005 (MicroChem) was spin coated, pre-baked (65 ℃ and 95 ℃ for 1 min and 4 min, respectively), shaped using a mask aligner, post-baked (65 °C and 95 °C for 1 min and 4 min, respectively) and developed in SU-8 developer (MicroChem). The SU-8 was hard-baked at 180 °C for 1 hour after development. Next, LOR 3A (MicroChem) was spin-coated and baked (180 °C, 10 min) as a lift off resist, on top of which a trace layer was shaped using S-1818 (MicroChem, 115 °C, 1 min). Prior to sputtering deposition, a short duration of oxygen plasma clean (SuperPlasmod 300) was applied (100 W, 30 s) to improve the adhesion. Then a layer of plain platinum was sputtered (25 W, 3 mTorr, 100 nm). We found that this process produced adequate adhesion without the need for a metallic adhesion layer. In fact, we found that metallic adhesion layers increased the stress in the films causing them to curl or “self-roll”^29^ which was undesirable for our applications. After deposition, the wafer was washed in acetone to lift-off unwanted platinum. Following deposition of the trace layer, the porous platinum layer was applied with sputter deposition, identical to the previous step except using a different pattern with higher pressure and power settings (100 W, 96 mTorr, 100 nm). The porous layer was applied such that every other electrode pad was coated, alternating porous and flat down the length of the electrode (figure 1(a)). Then, a 5-μm-thick top insulation layer of SU-8 2005 was patterned through the same fabrication process as fabricating the first SU8 layer. The wafer was then hard-baked for 1 hour at 180 °C. After cooling, the wafer was put into MF321 (MicroChem) to etch the aluminum release layer. After roughly 8 hours, electrodes were transferred to distilled water, where they remained for a minimum of 48 hours before future handling to ensure MF321 was washed off. After washing, electrodes were mounted to custom PCB breakout boards (OSH Park) using silver epoxy (Silver Print II, GC Electronics).

It is important to note that here we use two metal deposition steps so that we can compare flat and porous Pt electrodes on the same device. If we were to fabricate a device with only porous Pt electrodes we could make such a device with a single metal deposition step like previously reported flexible electrodes^5^.

### 2.2 Benchtop testing of electrodes

We performed electrochemical impedance spectroscopy (EIS) and cyclic voltammetry (CV) on each electrode using a Gamry Potentiostat (the Reference 600+). In both EIS and CV we used a 3-electrode setup to avoid any artifacts due to the impedance of the counter electrode^30^, with the working electrode (WE) and working sensing (WS) leads both connected to the experimental electrode, the counter electrode (CE) connected to a 99.9% platinum wire, 0.010” in diameter (uGems), and the reference (RE) and reference sensing (RS) electrodes connected to an encapsulated Ag/AgCl wire (Gamry Instruments). EIS was measured from 100 kHz to 1 Hz with 30 points per decade at a voltage of 10 mV against the reference electrode. Cyclic voltammetry was performed from −0.6 to 0.8 V at a scan rate of 100 mV/sec. 8 full scans were performed and cathodic charge storage capacity (cCSC) was calculated by first taking the integral of the two curves where each was below 0 A, subtracting one integral from the other and taking the absolute value. This is essentially the same as taking the area between the two curves where they were both under 0 A.

We also performed background thermal noise measurements of each electrode in 1X phosphate-buffered saline (PBS) (Research Products International 10X diluted) by connecting our custom break-out board to an RHD2132 amplifier (Intan Technologies) and an OpenEphys acquisition board (OpenEphys). The typical RMS noise of the Intan amplifier is 2.4 μV^31^. As a reference electrode, a stainless-steel surgical screw was submerged in the same solution. A surgical screw was chosen because this is the same kind of reference that would be used in an in vivo setting. Signals were viewed and recorded via the open-source RHD2000 GUI interface (Intan). Signal extraction and filtering were performed using Python 3 or MATLAB (Mathworks). Saline noise recordings were bandpass filtered from 300-5000 Hz, the same range used to filter our in vivo spike data.

### 2.3 Acute in vivo recordings from electrodes

In order to overcome the buckling force that the brain applies to flexible electrodes, they require a shuttle^5,32,33^, stiffener^34^, or tension^35^ in order to be implanted. Here we used a silicon shuttle to implant the electrodes using similar methods and materials found in previously published literature^32^. Before implantation, all electrodes were bound to silicon shuttles (stiffeners) roughly 50 μm thick by 100 μm wide using a thin layer of polyethylene glycol (PEG) between the electrode and stiffener. Care was taken to ensure that electrodes were facing outward and away from the stiffener.

All rodent work was approved and performed under the guidelines of the Rice University Institutional Animal Care and Use Committee. We used adult male Long Evans rats between 300 and 600 grams. Rats were induced with 5% isoflurane in oxygen and maintained under 1-2.5% isoflurane as needed. Prior to the start of surgery, rats were dosed with 0.03 mg/kg buprenorphine subcutaneously and 2% lidocaine subcutaneously at the scalp. Lidocaine cream was also applied to the ears to minimize discomfort from ear bars. After opening the scalp and cleaning the skull, two screws were placed in the skull. The first was placed over left frontal bone to act as an electroencephalogram (EEG), and the second was placed over cerebellar bone to act as an electrical reference for the EEG and implanted electrode. Next, a craniotomy was made over the right frontal bone and dura was carefully removed from the brain. Next, the electrode/stiffener combination was lowered into the brain at a rate of about 200 μm/sec, aiming for motor cortex (coordinates: +1.8 A/P, +2.5 M/L, −1.9 D/V). Small adjustments in A/P and M/L coordinates were made to avoid large blood vessels whenever necessary. After implantation, we waited a minimum of 1 hour before recording in order to allow tissue to settle and brain activity to normalize. Isoflurane levels were also adjusted as needed to maintain a good ratio of burst-suppression activity. The brain was kept well irrigated with saline to ensure it did not dry out. After recording, rats were euthanized with an injection of Euthasol.

Aside from the electrode placements as detailed above, recordings were performed using the same equipment as in benchtop testing. EEG signals were bandpass filtered from 10 to 50 Hz to maximize the burst-suppression signal for discrimination between bursts and suppressions. Electrode signals were bandpass filtered from 300-5000 Hz to isolate spikes.

After the in vivo recording was complete, the electrodes were removed from the brain and soaked in distilled water to wash off any remaining PEG, then subjected to another round of benchtop testing as described above.

## 3. Results

### 3.1 Benchtop testing of electrodes

We created two different designs of 16-channel flexible electrodes using SU-8 and platinum with square electrode pads that were 100 μm^2^ or 400 μm^2^. The electrodes were designed such that porous and flat electrode pads alternated along the length of the electrode shank (figure 1(a)). We subjected each electrode to EIS and CV to characterize their electrical properties (figure 2(a)-(d)). We compared the impedance of porous and flat electrodes to each other at 1 kHz, because most neural activity is especially strong at this frequency and it is well established for comparing impedance values reported in literature.

**Figure 2:**
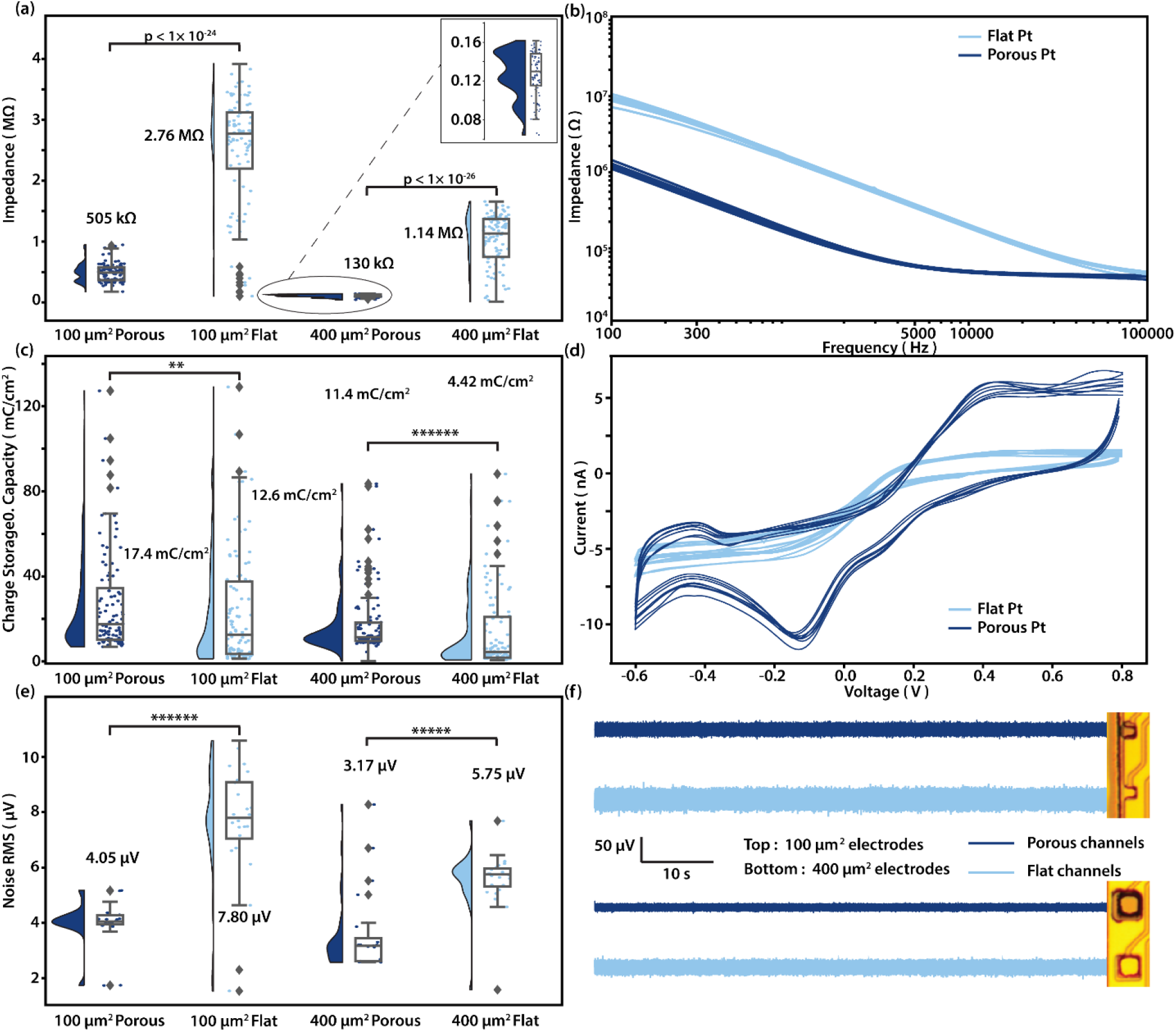
Pre- implantation electrical characterization and noise recording in PBS of porous and flat platinum electrodes. **(a)** Raincloud plot shows impedance at 1 kHz of 100 μm^2^ porous and flat electrodes, and 400 μm^2^ porous and flat electrodes, respectively. The y-value for each scatter dot represents the impedance measured for one electrode of the type described in corresponding to the x-axis label. The x-position of the dots are randomly jittered to aid visualization. Numbers shown are medians. (n = 100, 97, 93, 92; One-sided Mann–Whitney U test). **(b)** EIS of all 16 channels in a 400 μm^2^ electrode. Porous channels are in dark blue while flat channels are in light blue. **(c)** Cathodic charge storage capacity calculated for 100 μm^2^ porous and flat electrodes, and 400 μm^2^ porous and flat electrodes, respectively. Numbers shown are medians. (n = 100, 101, 94, 95; **p < 0.01, ******p < 0.000001; One-sided Mann–Whitney U test). **(d)** Current over voltage curves of all 16 channels in a 400 μm^2^ electrode for a single run of cyclic voltammetry from −0.6 to 0.8 V. Porous channels are in dark blue while flat channels are in light blue. **(e)** Root mean square(RMS) of noise recorded in PBS for 100 μm^2^ porous and flat electrodes, and 400 μm^2^ porous and flat electrodes, respectively. Numbers shown are medians. (n = 24, 24, 24, 24; ******p < 0.000001, *****p < 0.00001; One-sided Mann–Whitney U test). **(f)** Traces of background noise measured in PBS from 100 μm^2^(Top) porous(Dark blue) and flat electrodes(Light blue), and 400 μm^2^(Bottom) porous(Dark blue) and flat electrodes(Light blue). Band-pass filtered from 300-5000 Hz.

For 400 μm^2^ electrodes, the median impedance for flat electrodes was 1.14 MΩ, while for porous electrodes it was 130 kΩ, constituting a 9-fold decrease. For 100 μm^2^ electrodes, flat electrodes median impedance was 2.76 MΩ and porous electrodes median impedance was 505 kΩ. We also calculated the specific impedance, which is defined as impedance times electrode area, as well as the cCSC. All the results of specific impedance and cCSC are shown in Table 1, in the format of median, 0.95 confidence interval (C.I.).

**Table 1.** Pre- implantation electrical characterization results in the format of median (0.95 confidence interval of the median).

| | 100<br>$\mu\text{m}^2$ Porous | 100 $\mu\text{m}^2$<br>Flat | 400 $\mu\text{m}^2$<br>Porous | 400 $\mu\text{m}^2$<br>Flat |
| --- | --- | --- | --- | --- |
| Impedance at 1 kHz (kΩ) | 505<br>(466-537) | 2760<br>(2630-2820) | 130<br>(123-139) | 1140<br>(1020-1250) |
| Specific Impedance<br>($\Omega\cdot\text{cm}^2$) | 0.505<br>(0.466-0.537) | 2.76<br>(2.63-2.82) | 0.520<br>(0.492-0.556) | 4.56<br>(4.08-5.00) |
| cCSC (mC/cm$^2$) | 17.4<br>(14.2 - 23.0) | 12.6<br>(6.32 - 20.7) | 11.4<br>(10.6-12.7) | 4.42<br>(3.08-7.10) |

As expected, the specific impedance of the porous electrodes is nearly the same for the 400 μm^2^ and 100 μm^2^ electrodes (~3%); however, we were surprised to find that for flat electrodes, the specific impedance is ~65 % higher for 400 μm^2^ compared to 100 μm^2^ electrodes. A likely explanation is that the impedance of an electrode consists of a surface impedance in parallel with an edge impedance. The total conductance is thus the sum of the surface conductance and the edge conductance. As electrodes get smaller, the surface conductance decreases as 1/r^2^, while the edge conductance decreases more slowly as 1/r (where r is the width of an electrode). As a result, the surface conductance will eventually become small compared to the edge conductance. At this point, the specific impedance (impedance times area) will be lower than predicted based on surface impedance. In other words the edges dominate the electrode impedance. This critical size will be larger for high impedance surfaces like flat Pt. As a result, the porous Pt impedance could remain dominated by surface impedances for similar sized electrodes.

We excluded from our data ~20 % of 100 μm^2^ electrodes and ~5 % of 400 μm^2^ electrodes because they were shorted or had badly connected traces as determined by our exclusion criteria. We defined “badly connected” channels with impedances that were more than 4 MΩ, which is 45 % larger than the highest median value of all 4 types of electrodes, and shorted channels as those flat channels with impedances less than 200 % of the adjacent porous channel. Of the electrodes that were not excluded due to shorted and bad connections, an average of 74% fell within ±30% of the median impedance value (figure S2). The variability of impedance values is likely due to process variation and may reflect alignment errors during the electrode metallization step.

### 3.2 Benchtop noise testing of electrodes

After measuring impedance, we next performed a recording with the electrode array in PBS and a reference screw in the same solution roughly 5 cm away. Median root mean square noise across 400 μm^2^ electrode pads was 3.17 (2.62-3.31) μV (median, 0.95 C.I.) for porous electrodes and 5.75 (5.36-5.96) μV (median, 0.95 C.I.) for flat electrodes. For 100 μm^2^ electrode pads, median RMS noise was 4.05 (3.95-4.28) μV (median, 0.95 C.I.) for porous electrodes and 7.80 (7.16-9.07) μV (median, 0.95 C.I.) for flat electrodes (figure 2(e)-(f)). Measuring in PBS, thermal noise should be the dominant component of the observed signal, so we next calculated the theoretical thermal noise for each electrode based off of the impedance measured earlier. The theoretical thermal noise values and match well with that of measured, the actual noise RMS varied from the theoretical value by 22.5 (9.95-24.8)% (median, 0.95 C.I.) (figure.S3).

### 3.3 In vivo testing of electrodes

Next, we sought to test whether decreased impedance within porous electrodes led to any notable improvements in *in vivo* signal amplitude. To this end, we recorded from motor cortex of rats under isoflurane anesthesia. Rodents under isoflurane anesthesia show burst-suppression cortical activity characterized by bursts of activity lasting about 1-3 seconds (bursts) interspersed by longer periods of silence (suppressions). Burst-suppression is an ideal condition in which to test electrode performance, as chewing and motion artifacts are non-existent during neural activity, and suppressions give clean examples as to the noise background performance of electrodes in vivo.

From recordings of burst-suppression activity in motor cortex (figure 3(a)), we were able to isolate neural activity and compare the noise RMS during suppression between porous and flat electrodes (figure 3(b)). In 100 μm^2^ electrodes, we found that median noise RMS from suppression recorded from porous electrodes was ranged from 7.29 (6.28-8.01) μV (median, 0.95 C.I.), while the median for flat electrodes was 8.75 (7.89-9.87) μV (median, 0.95 C.I.). And that of 400 μm^2^ electrodes are 5.11 (4.81-5.65) μV (median, 0.95 C.I.) and 5.91 (5.63-7.55) μV (median, 0.95 C.I.) for porous and flat electrodes, respectively.

**Figure 3:**
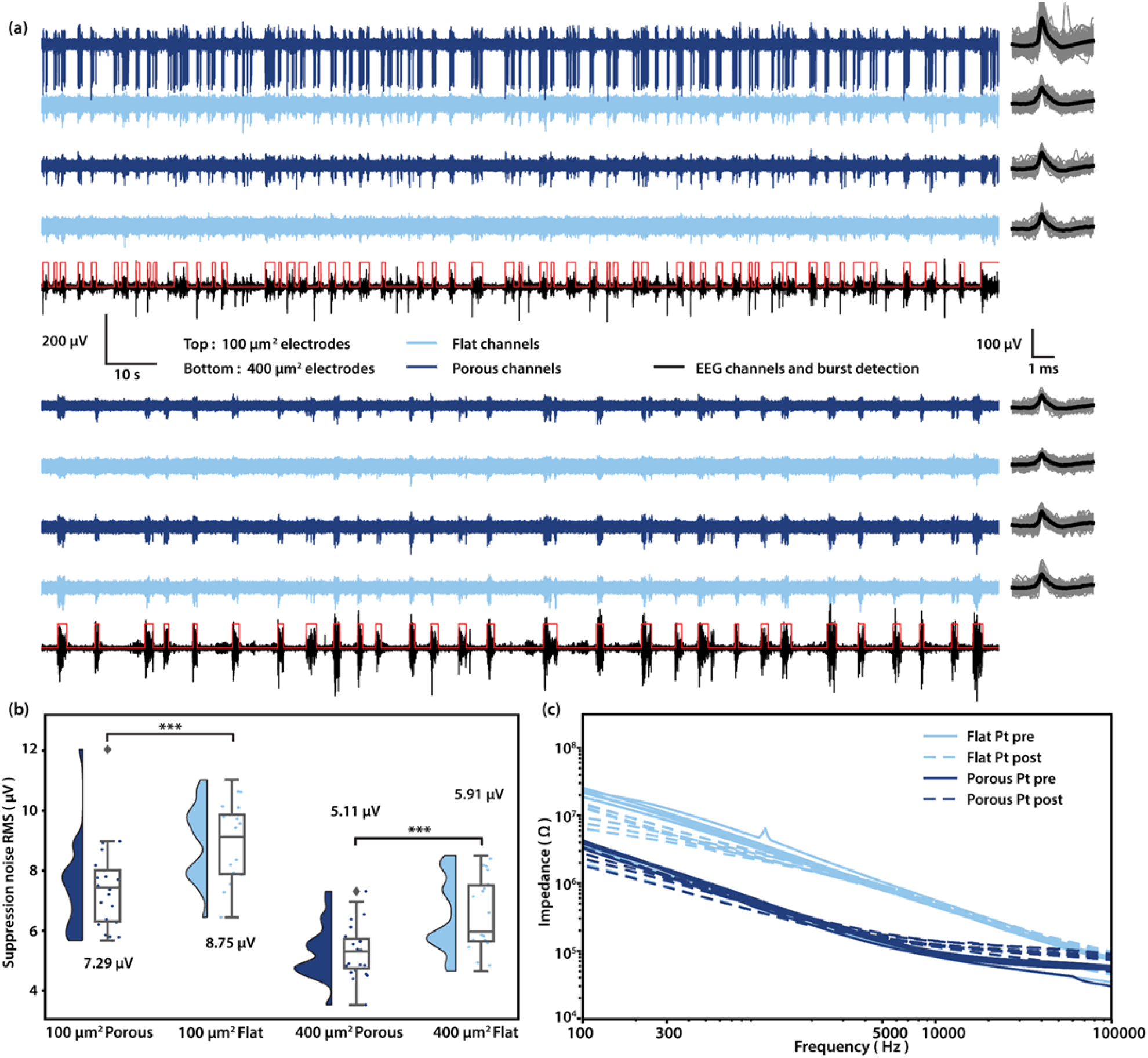
Acute in vivo performance and pre-/post- implantation EIS of porous and flat platinum electrodes. **A** Recorded traces and mean waveform of 100 μm^2^ (Top) porous (Dark blue) and flat electrodes (Light blue), and 400 μm^2^ (Bottom) porous (Dark blue) and flat electrodes (Light blue). Spike waveforms (Gray) are extracted and plotted on the right, mean waveforms are shown in black. The traces in black are EEG channels, which are used to determine burst (Highlighted in red) and suppression periods. **B** Noise RMS during suppression of 100 μm^2^ porous and flat electrodes, and 400 μm^2^ porous and flat electrodes, respectively. Numbers shown are medians. (N = 22, 21, 23, 22. ***p < 0.001, ***p < 0.001; One-sided Mann–Whitney U test). **C** Pre- and post- implantation EIS of all 16 channels in a 400 μm^2^ probe.

After our recording tests were complete, we removed the electrode from the rat brain and re-tested the electrical properties using EIS (figure 3(c)). When comparing pre and post-implant values for impedance, we could see that the porous electrodes continued to show significantly reduced impedances compared to flat Pt suggesting that directly sputtered platinum electrodes are durable enough to undergo significant handling without damage to the porous platinum layer. Future work is needed to determine if the good mechanical stability we observed in our experiments translates to stable chronic performance in vivo.

### 3.4 Electrode geometry affects impedance reduction

In section 3.1, we proposed that both surface impedance and edge impedance contribute to the total impedance, which suggests that electrode geometry could determine impedance. To illustrate how electrode geometry can affect the impedance, we created flat flexible probes with three differently-shaped electrodes (figure 4(c)) that we refer to as: square, ring, and waffle designs. All three have the same footprint (3025 μm^2^). The ring and waffle electrodes were designed such that they have identical total surface area (2125 μm^2^), but different perimeters (340 μm and 700 μm, respectively). We performed EIS as before. We see from electrochemical impedance spectroscopy that at 1 kHz, all three electrode shapes are well below the cutoff frequency of the Randles cell (figure. S5). Thus, the 1 kHz impedance values represent the electrode surface impedance for all three electrode geometries. Interestingly, we found that while the square electrodes have a surface area 1.42 times of that of the ring electrode, their impedance was not significantly lower as we would expect if we neglected edge effects (figure 4(a)). After normalizing by the surface area, the square electrodes show a specific impedance that is significantly higher compared to waffle electrodes(p < 0.01, figure 4(b)). These data suggest that increasing surface area does not always reduce impedance, and that increasing electrode edges is an alternative route toward reduced electrode impedance.

**Figure 4:**
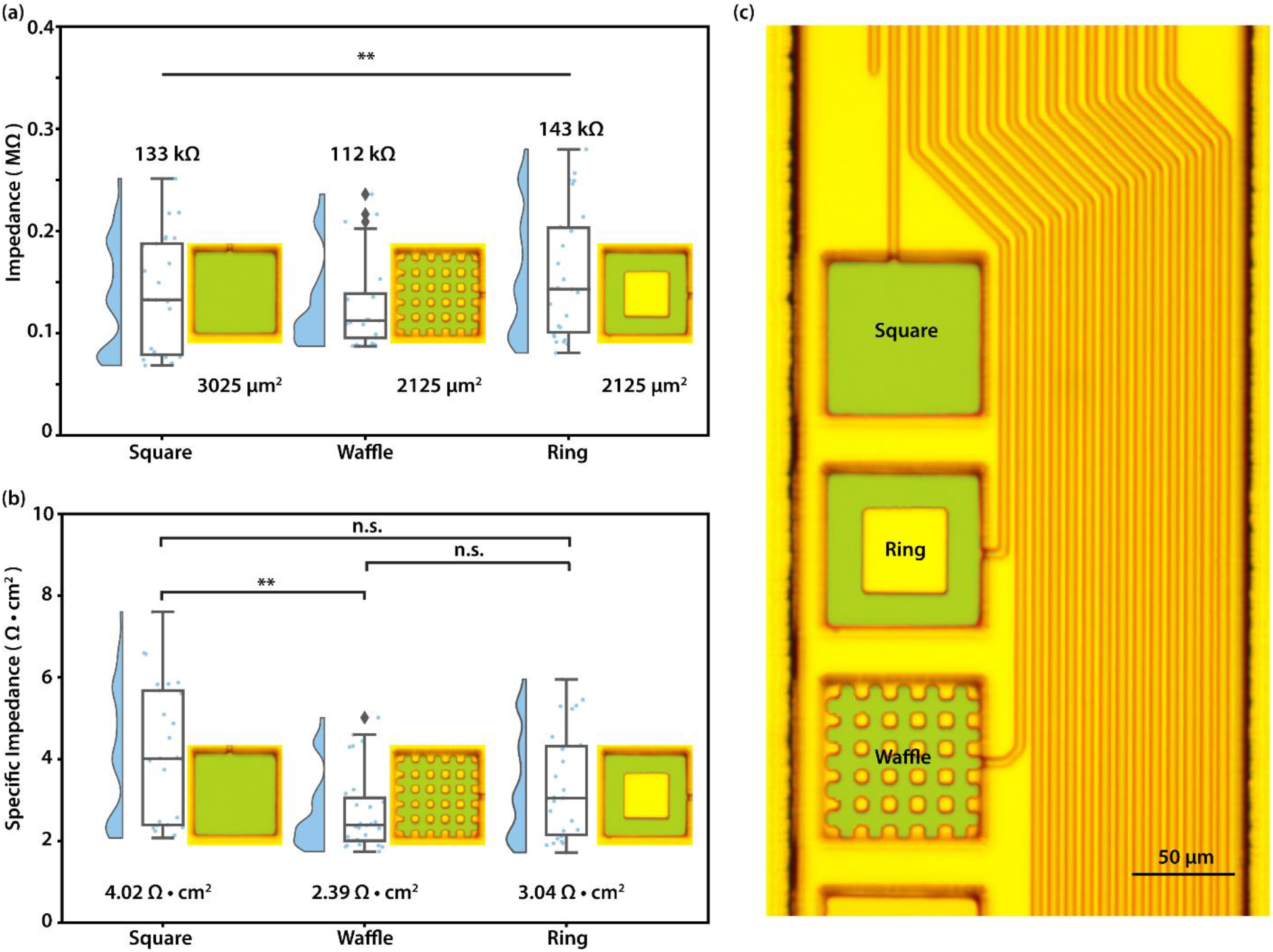
Impedance characterization shows effects of electrode geometry. **(a)** Collected impedance at 1 kHz of ring, waffle and square electrodes. n = 25, 27, 26; **p < 0.01; Kruskal-Wallis H Test. **(b)** Specific impedance at 1 kHz of ring, waffle and square electrodes(n = 25, 27, 26, **p < 0.01; [waffle,square]; Kruskal-Wallis One-way ANOVA with a post-hoc Dunn’s multiple comparison test). **(c)** False-colored ring, waffle and square flat platinum electrodes (Green: Flat Pt). Scale bar 50 um.

## 4. Conclusion and discussion

Here, we demonstrated a new method for direct sputtering of a porous platinum electrode that shows a 5-9 fold impedance reduction compared to flat platinum. Direct sputtering has the advantage of being able to apply an impedance-reducing coating to the entire wafer at once, eliminating the need to electrically connect or handle individual electrode arrays. Thus this method easily scales to high-channel-count probes, regardless of how many electrodes are placed on a single implant. Our results show that we are able to reliably decrease the impedance of electrodes simply by using an existing metallization step with different deposition settings. While direct sputtering does significantly lower the impedance of our electrodes, our impedance reduction ratio is not as high as what is more commonly seen with PEDOT^13^. However, a rougher porous Pt coating can be achieved with more aggressive sputtering settings, which requires a sputtering system with greater tolerance to higher power and higher pressure.

One finding of note is that change in specific impedance scales differently for 400 μm^2^ electrodes versus 100 μm^2^ electrodes, and for porous versus flat. This data implies that for small electrodes, the edges may play an important role in the overall electrode impedance. The influence of edges is further highlighted by the fact that electrodes with a large edge to surface ratios like ring or waffle shaped electrodes have a lower specific impedance compared to square electrodes (figure 4(b)). Note that because the electrode layer is only 100 nm thick, the electrode sidewalls do not significantly contribute to the total surface area. Concentration of field lines near the edges of neural electrodes are known to play a role in the safety and efficacy of neural stimulation electrodes^36^ and changing the perimeter or electrode shape can alter the electrical properties of these stimulation^37,38^. While the most profound effects are on current density for stimulation, changing the shape of an electrode has been reported to also affect impedance. In one case, creating more curves and edges on a fractal electrode did not significantly affect impedance, while in another, creating a segmented electrode decreased impedance when the total area of a large electrode was split into several segments. One possible explanation is that there is a characteristic length where edge effects are most pronounced. Similar effects may explain the differences in impedance that we observe, but it is not clear what the consequences will be for neural recording. Future work will be needed to answer these questions.

There has recently been debate as to what extent impedance reduction is useful to improve neural recordings^39,40^. In fact, in the case of intracellular recordings, raising the impedance from the standard 5 MΩ to several hundred MΩ still produced good intracellular measurements provided that one uses appropriate amplifying electronics^41^. Recent publications in a variety of materials have focused on this same idea of impedance reduction being a key measure of a good electrode, without explaining how this impedance value will affect the quality of the recording. Large recording platforms like neuropixels^7^ or flexible multi-shank systems^8^ have impedances of around 1 MΩ. In theory, reducing impedance is critical for quality neural recording: on one hand, as seen in our data, low-impedance electrodes have reduced noise. Because thermal noise is proportional to the square root of electrode impedance, there are diminishing returns on thermal noise reduction as one continues to reduce the impedance. On the other hand, the recording electrodes and the amplifier create a voltage divider, which can attenuate the signal if the electrode impedance is comparable to the input impedance of the amplifier or the stray impedance to ground. Typical value of the input impedance of the amplifier is around 10-13 MΩ^10,39^. High impedance electrodes will greatly reduce the recording signal, for example, a 10 MΩ electrode will attenuate the recording signal to ~50%. Thus, many publications suggest that it is not necessary to prioritize low impedance once it is significantly below the input impedance of the amplifier. Neto et al. (2018)^39^ claimed that the electrodes within a range of 0.1-2 MΩ should not have a great impact on signal. Ludwig et al. (2011)^42^ made a similar statement that for impedances near 5 MΩ, impedance reduction improves recording quality. Total noise gathered by an electrode is equal to the root sum of the squares of each of the individual noise sources, so the greatest source of noise will tend to dominate the noise floor. Proximity to the source of neural signal plays a large role in the size of detected spikes^6^, meaning that while low impedance can be permissive in acquiring quality neural recordings, having sufficient coverage and density is often more important once the thermal noise due to electrode impedance falls below other noise sources (like background biological noise). In addition, from where the electrode is recording from the neuron can be a game changer. Bakkum et al. (2019)^43^ has shown that signal amplitude reach millivolt levels for small electrodes near the axon initial segment (AIS), while near the end of axon and dendrites it drops to less than 100 μV. If impedance reduction is no longer the largest obstacle in advancing extracellular recording electrodes, what other parameters should be considered?

One often undervalued parameter is the cost and ease of manufacturing. Though many varieties of experimental electrode have been published in the last decade, relatively few designs have proliferated and become commonly used. Outside of engineering journals, it is much more common to see decade-old Neuronexus or TDT electrodes used for neuroscience research instead of newer, higher-channel-count electrodes. One possible explanation for this could be challenges in scaling up manufacturing from small experimental batches to large production runs. Neuropixels^7^ is one example of a new widely-available commercial electrode, and also relies on a deposited electrode material followed by a hydrothermal treatment^25^.

Another important consideration is electrode size and density. Large electrodes have lower impedance and they have a greater chance to record from neurons. However, they result in attenuated signals from single-unit because the high amplitude signal from a local neuron will be averaged with other smaller signals^44^, which is usually considered as background biological noise. In the case of cells cultured on microelectrode arrays, this attenuation can be described as a shunt impedance that arises from the area of the electrode that is not covered by the cell^45^. Single-unit recording requires small and high-density electrodes, because they allow for a higher chance to pick up a neuron without significant spatial averaging. The porous Pt film we reported here provide a reliable and simple way to reduce impedance for smaller electrodes. Another issue when the electrodes become smaller is that they are more difficult to fabricate due to the resolution limit of photolithography and an alternative E-beam lithography will greatly increase the cost of fabrication.

Overall our results show that porous Pt is an effective method to create robust, low-impedance electrodes on flexible substrates that is compatible with wafer-scale manufacturing, and increasing edges of the electrodes also help reduce the impedance; however, when designing neural electrodes, a number of factors will affect the overall impedance values and ability of electrodes to isolate individual action potentials depends on many factors in addition to the impedance value at 1 kHz.

## Acknowledgment

We thank Ben Avants for assistance with electronics for in vivo recording, Rice Shared Equipment Authority for the support in fabrication. This work was funded by NIH grant R21EY028397A and the Dan Duncan Postdoctoral Fellowship(AVR).

