## Supplementary Information for "Sputtered porous Pt for wafer-scale manufacture of low-impedance flexible microelectrodes"

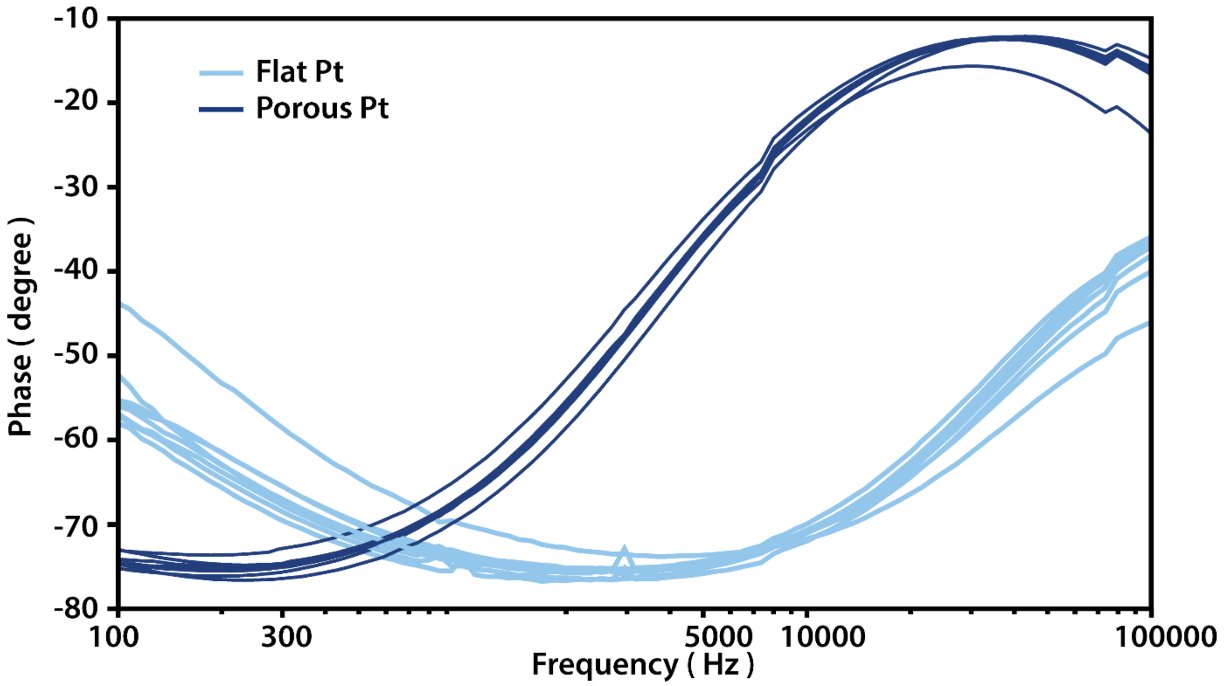


**Figure S1**. Phase plot of all 16 channels in a 400 µm$^2$ electrode.

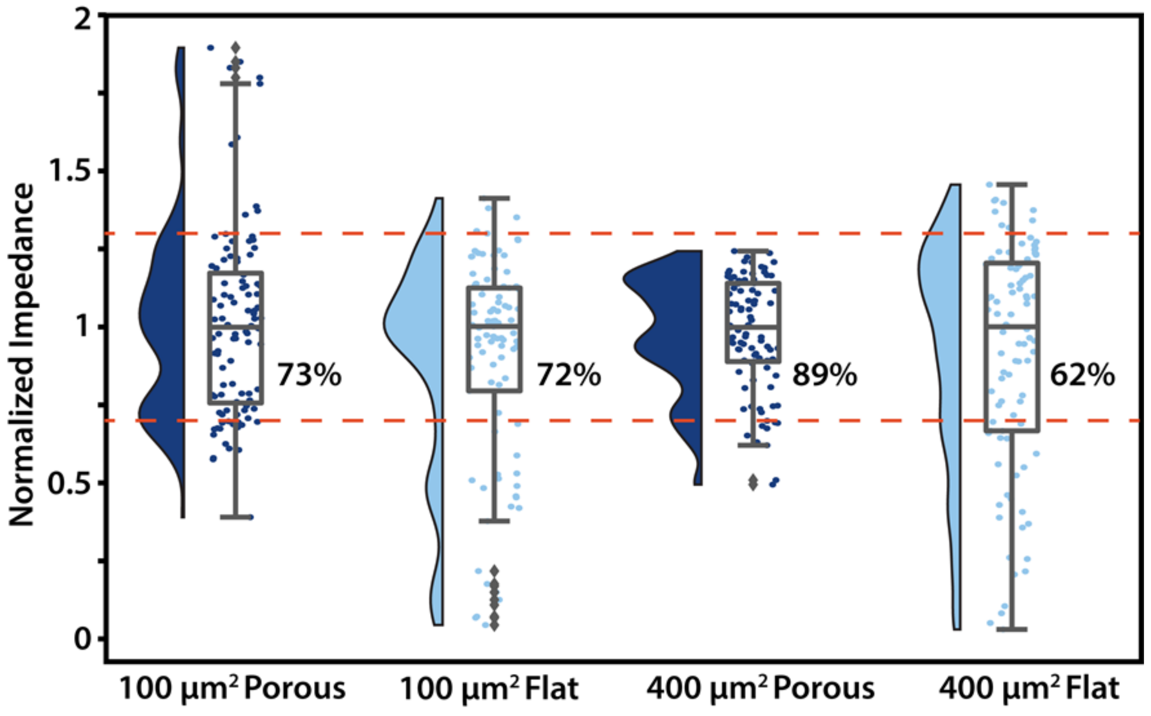


**Figure S2.** Normalized impedance and the percentage of the electrodes fell within ±30% of the median value.

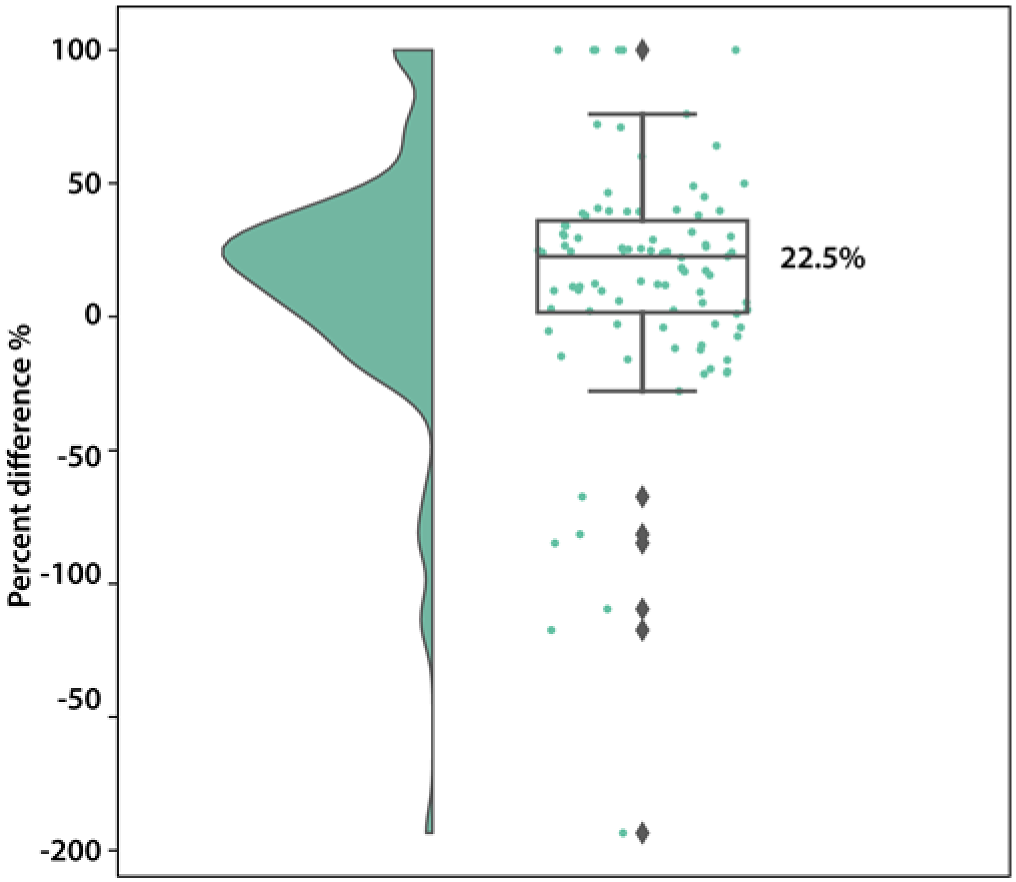


**Figure S3:** Raincloud plot shows the percent difference between recorded RMS of noise in PBS and theoretical thermal noise. Median is shown. (n = 95).

The theoretical voltage RMS is calculated using the following equation:

$$V_{RMS} = \sqrt{\int_{f_{c1}}^{f_{c2}} 4 \cdot k_B \cdot T \cdot Z_{real} df}$$

Where $f_{c1}$ and $f_{c2}$ are the cut-off frequencies, which are 300 Hz and 5000 Hz in this work. $k_B$ is the Boltzmann constant, T the temperature and $Z_{real}$ the real part of the impedance.

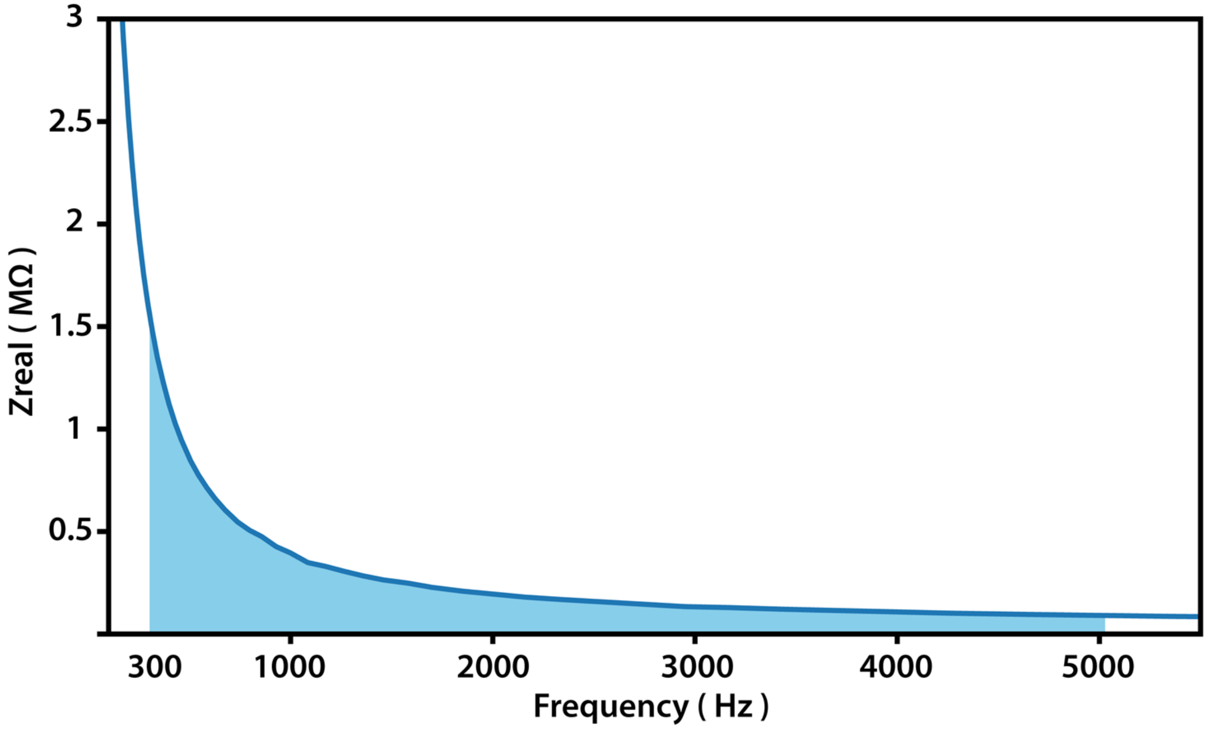


**Figure S4.** The integral area used to calculate the theoretical thermal noise.

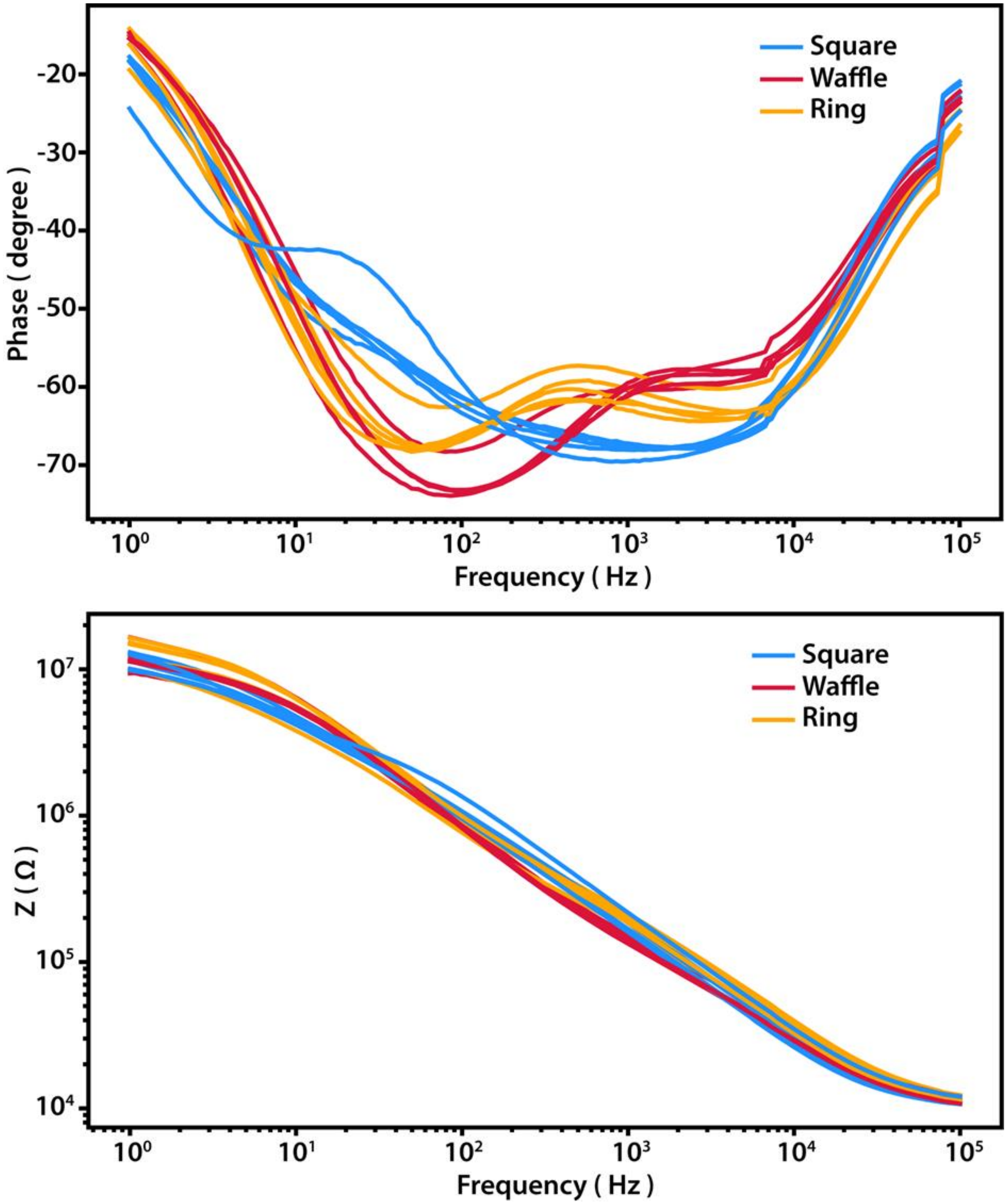


**Figure S5.** Bode plot of square, waffle and ring electrodes from a probe.
